# Outcomes of the 2019 EMDataResource model challenge: validation of cryo-EM models at near-atomic resolution

**DOI:** 10.1101/2020.06.12.147033

**Authors:** Catherine L. Lawson, Andriy Kryshtafovych, Paul D. Adams, Pavel V. Afonine, Matthew L. Baker, Benjamin A. Barad, Paul Bond, Tom Burnley, Renzhi Cao, Jianlin Cheng, Grzegorz Chojnowski, Kevin Cowtan, Ken A. Dill, Frank DiMaio, Daniel P. Farrell, James S. Fraser, Mark A. Herzik, Soon Wen Hoh, Jie Hou, Li-Wei Hung, Maxim Igaev, Agnel P. Joseph, Daisuke Kihara, Dilip Kumar, Sumit Mittal, Bohdan Monastyrskyy, Mateusz Olek, Colin M. Palmer, Ardan Patwardhan, Alberto Perez, Jonas Pfab, Grigore D. Pintilie, Jane S. Richardson, Peter B. Rosenthal, Daipayan Sarkar, Luisa U. Schäfer, Michael F. Schmid, Gunnar F. Schröder, Mrinal Shekhar, Dong Si, Abishek Singharoy, Genki Terashi, Thomas C. Terwilliger, Andrea Vaiana, Liguo Wang, Zhe Wang, Stephanie A. Wankowicz, Christopher J. Williams, Martyn Winn, Tianqi Wu, Xiaodi Yu, Kaiming Zhang, Helen M. Berman, Wah Chiu

## Abstract

This paper describes outcomes of the 2019 Cryo-EM Map-based Model Metrics Challenge sponsored by EMDataResource (www.emdataresource.org). The goals of this challenge were (1) to assess the quality of models that can be produced using current modeling software, (2) to check the reproducibility of modeling results from different software developers and users, and (3) compare the performance of current metrics used for evaluation of models. The focus was on near-atomic resolution maps with an innovative twist: three of four target maps formed a resolution series (1.8 to 3.1 Å) from the same specimen and imaging experiment. Tools developed in previous challenges were expanded for managing, visualizing and analyzing the 63 submitted coordinate models, and several novel metrics were introduced. The results permit specific recommendations to be made about validating near-atomic cryo-EM structures both in the context of individual laboratory experiments and holdings of structure data archives such as the Protein Data Bank. Our findings demonstrate the relatively high accuracy and reproducibility of cryo-EM models derived from these benchmark maps by 13 participating teams, representing both widely used and novel modeling approaches. We also evaluate the pros and cons of the commonly used metrics to assess model quality and recommend the adoption of multiple scoring parameters to provide full and objective annotation and assessment of the model, reflective of the observed density in the cryo-EM map.

## Introduction

Electron cryo-microscopy (cryo-EM) has emerged as a key method to visualize and model a wide variety of biologically important macromolecules and cellular machines. Researchers can now routinely produce structures at near-atomic resolution, yielding new mechanistic insights into cellular processes and providing support for drug discovery^1-3^. Many academic institutions and pharmaceutical companies have invested in modern cryo-EM facilities, and multi-user resources are opening up worldwide^4^.

The recent explosion of cryo-EM structures raises important questions. What are the limits of interpretability given the quality of the maps and resulting models? How do we quantify model accuracy and reliability under the simultaneous constraints of map density and chemical rules?

The EMDataResource Project (EMDR) was formed in 2006 as a collaboration between scientists in the UK (EMDataBank at the European Bioinformatics Institute) and the US (the Research Collaboratory for Structural Bioinformatics and the National Center for Macromolecular Imaging). Part of EMDR’s mission is to derive validation methods and standards for cryo-EM maps and models through community consensus^5^. We created an EM Validation Task Force^6^ analogous to those derived for X-ray crystallographic and NMR structures^7,8^ and have sponsored Challenges, workshops and virtual conferences to engage cryo-EM experts, modellers, and end-users^5,9-13^. During this period, cryo-EM has evolved rapidly (Figure 1).

**Figure 1.**
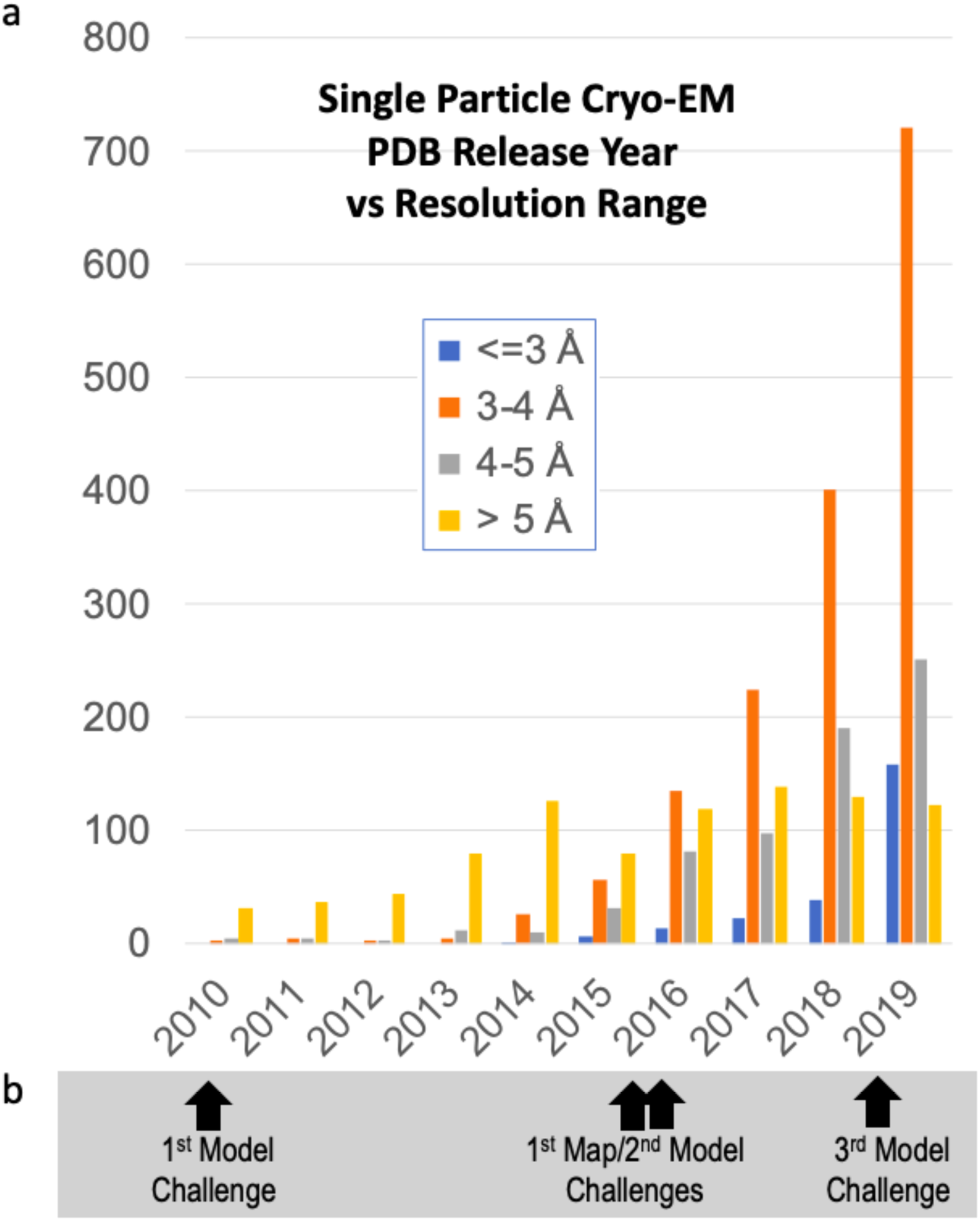
Single particle cryo-EM models in PDB. The availability of models in the Protein Data Bank^59^ derived from single particle cryo-EM maps has increased dramatically since the “resolution revolution” circa 2014^60^. (a) Plot shows that the steepest increase is for structures with reported resolution in the 3-4 Å range (orange bars). Higher resolution structures (blue bars), the topic of the Challenge presented here, are also beginning to trend upward. (b) EMDataResource Challenge activities timeline.

This paper describes outcomes of EMDR’s most recent Challenge, the 2019 Model “Metrics” Challenge. The goals were three-fold: (1) to assess the quality of models that can be produced using established as well as newly implemented modeling software, (2) to check the reproducibility of modeling results from different software developers and users, and (3) to compare the performance of model evaluation metrics, particularly fit-to-map metrics. Map targets were selected in the near-atomic resolution regime (1.8-3.1 Å) with an innovative twist: three form a resolution series from the same specimen/imaging experiment (Figure 2). The results lead to several specific recommendations for validating near-atomic cryo-EM structures directed towards both individual researchers and the Protein Data Bank (PDB) structure data archive.

**Figure 2.**
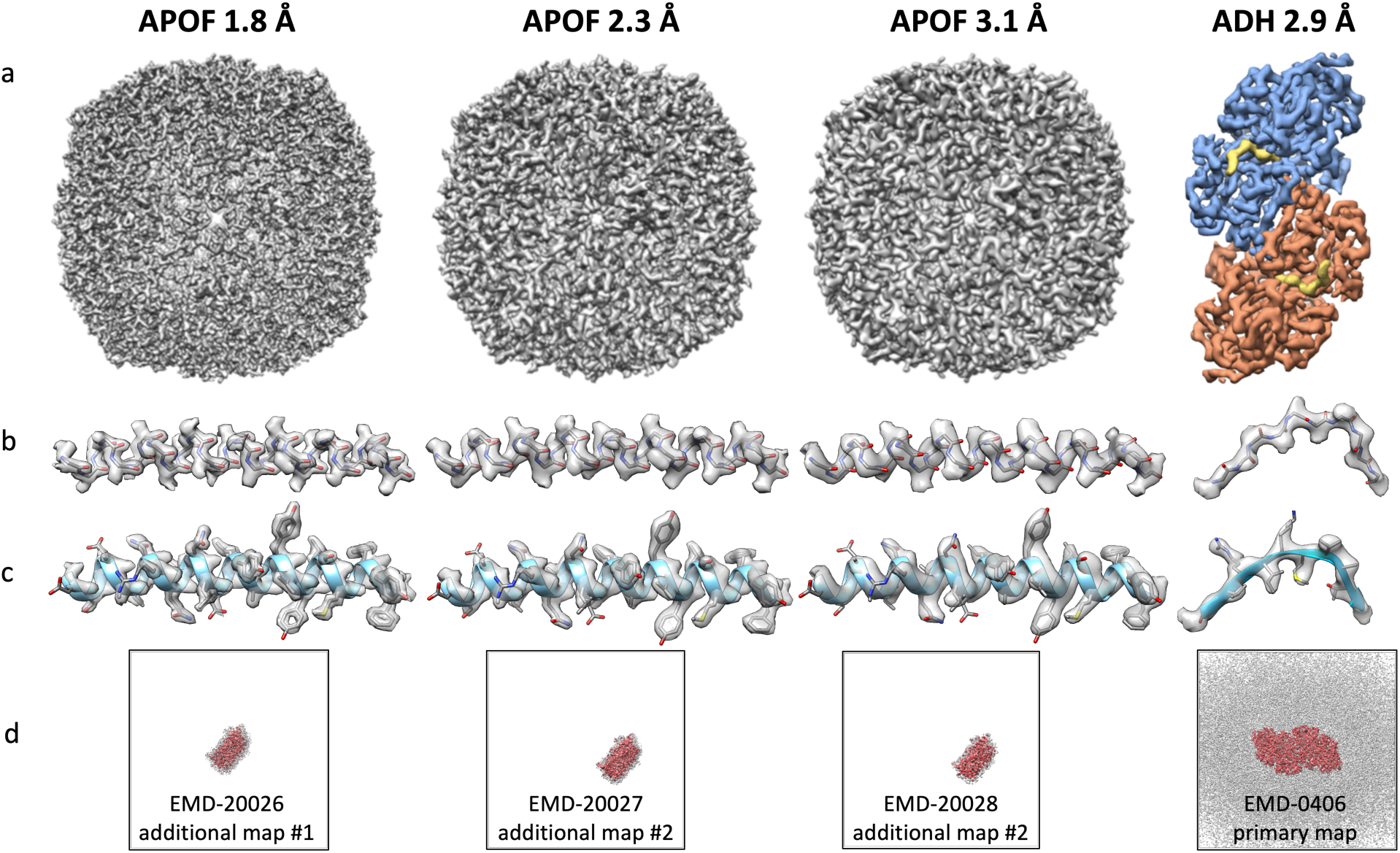
Challenge targets: cryo-EM maps at near-atomic resolution. Three density maps of α-helix rich apoferritin (APOF) at 1.8, 2.3, and 3.1 Å form a resolution series differing only in the number of particles averaged (EM Data Bank entries EMD-20026, EMD-20027, EMD-20028). The fourth density map, alcohol dehydrogenase (ADH) at 2.9 Å, contains both α-helices and β-sheets, as well as ligands (yellow density bound to blue and red subunits is NAD; EMDB entry EMD-0406). (a) Full map for each target. (b,c) Representative secondary structural elements (APOF: residues 14-42; ADH: residues 34-45) with masked density for protein backbone atoms only (b), and for all protein atoms (c). Over the 1.8-3.1 Å resolution range represented by the four target maps, visible map features transition from near-atomic to secondary structure-dominated. At 1.8 Å (APOF), most protein atom positions are well defined by the map density: backbone protrusions delineate carbonyl oxygen positions and holes appear inside aromatic rings. At 2.3 Å (APOF), most protein atom positions are within the contoured map density; carbonyl oxygen atom “bumps” in the map help to define direction of backbone trace. At 2.9 Å (ADH) and 3.1 Å (APOF), secondary structure features begin to predominate. Notably, carbonyl oxygen atoms “bumps” are absent in the map, making it harder to identify the direction of backbone trace. (d) EMDB maps used in model Fit-to-Map analysis (APOF targets: masked single subunits; ADH: unmasked sharpened map). The molecular boundary is shown in red (EMDB recommended contour level), background noise is represented in grey (1/3^rd^ of EMDB recommended contour level), and the full map extent is indicated by the black outline.

## Results

We describe here the pipeline and outcomes of the EMDR 2019 Model Metrics Challenge (Figure 3). Four maps representing the state-of-the-art in cryo-EM single particle reconstruction were selected as the Challenge targets (Figures 2, 3a). Three maps of human heavy-chain apoferritin (APOF), a 500 kDa octahedral complex of 24 α-helix-rich subunits, formed a resolution series differing only in the number of particles used in reconstruction (EMDB entries <u>EMD-20026</u>, <u>EMD-</u> <u>20027</u>, <u>EMD-20028</u>)^14^. The fourth map was horse liver alcohol dehydrogenase (ADH), an 80 kDa α/β homodimer with NAD and Zn ligands (<u>EMD-0406</u>)^15^.

**Figure 3.**
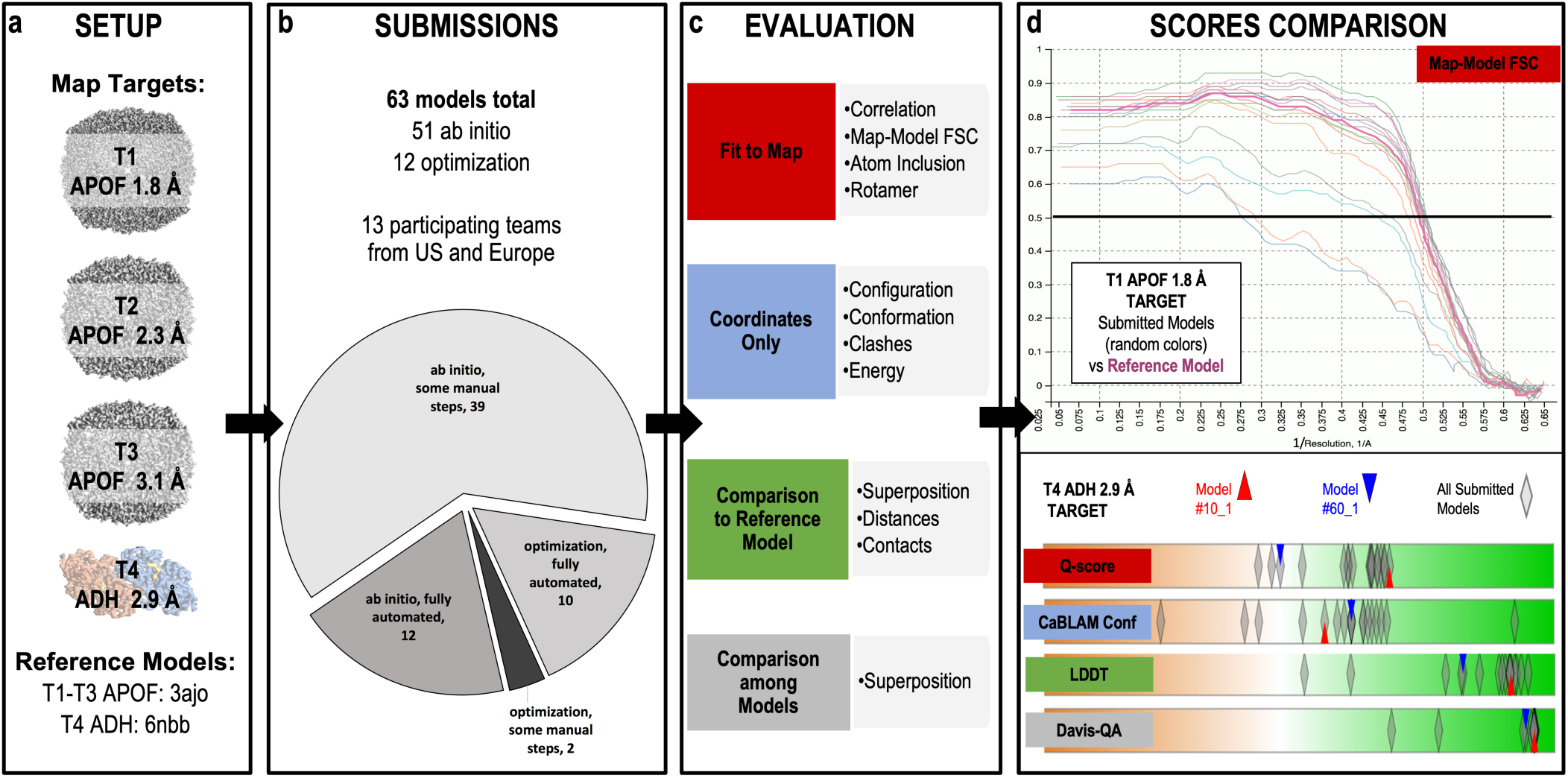
Challenge pipeline. *Setup* (a): A panel of experts selected four target maps in the 1.8-3.1 Å resolution range as well as reference models. *Submissions* (b): Methods involving *ab initio* and optimization, both with and without additional manual steps, were represented by 63 models submitted by 16 modeling teams. *Evaluation* (c): Building on the previous model challenge round^12^, the evaluation system was organized in four tracks (coordinates only, fit-to-map, comparison to reference model, and comparison among models), each with its own set of software tools for generating scores. *Scores Comparison* (d): Multiple interactive tabular and graphical displays enable comparative evaluations on the model-compare website (<u>model-compare.emdataresource.org</u>). *Top*: Map-Model Fourier Shell Correlation (FSC) curves for models submitted against the APOF 1.8 Å target (random light colors), versus curve for the reference model (bold cherry red). Map-Model FSC measures the overall agreement of the experimental density map with a density map derived from the coordinate model (model map)^17^. Curves are calculated from the Fourier coefficients of the two maps and plotted vs. frequency (resolution^-1^). Visualization of the full curve is useful to ensure that it follows the expected sigmoidal shape with the tail decaying exponentially. The resolution value corresponding to FSC=0.5 (black horizontal line) is typically reported, with smaller values indicating better fit. In this example, some overlaid curves indicate poor agreement between submitted model and the APOF 1.8 Å experimental map, but the majority indicate equivalence to or even improvement over the X-ray reference model (PDB entry 3ajo was rigid-body fitted to the cryoEM target map without further refinement). *Bottom:* scores comparison tool for ADH 2.9 Å target models. Interactive score distribution sliders reveal at a glance how well the submitted models performed relative to each other. Parallel lanes display score distributions for each evaluated metric. They are conceptually similar to the graphical display for key metrics used in wwPDB validation reports^7,38^. Displayed here are the score distributions for all models submitted against the ADH target for four representative metrics, one from each evaluation track, and scores for two individual models are also highlighted. Model scores are plotted horizontally (semi-transparent diamonds) with color-coding to indicate worse (left, orange) and better (right, green) values. Darker, opaque diamonds indicate multiple overlapping scores. The interactive display enables scores for individual models to be identified and compared (red and blue triangles). The model indicated by red triangles scored better than the model indicated by blue triangles for Fit-to-Map: Q-score, Comparison-to-Reference: LDDT, and Comparison-among-Models: Davis-QA, but the same model scored lower for Coordinates-only: CaBLAM Conformation. The example demonstrates the importance of evaluating models in each track: a high score in any one metric/track does not guarantee full optimization as judged by another metric.

A key criterion of target selection was availability of high quality experimentally determined model coordinates to serve as references. A 1.5 Å X-ray structure^16^ (PDB id <u>3ajo</u>) served as the reference for all three APOF maps, since no cryoEM-based model was available at the time. The X-ray model provides an excellent fit to each map, though not a fully optimized fit, owing to method/sample differences. The ADH reference was the model deposited by the original authors of the cryo-EM study (PDB id <u>6nbb</u>)^15^.

Thirteen teams from the US and Europe submitted 63 models in total, yielding 15-17 submissions per target (Figure 3b, Table I). The vast majority (51) were created *ab initio*, sometimes supported by additional manual steps, while others (12) were optimizations of publicly available models.

**Table I.**
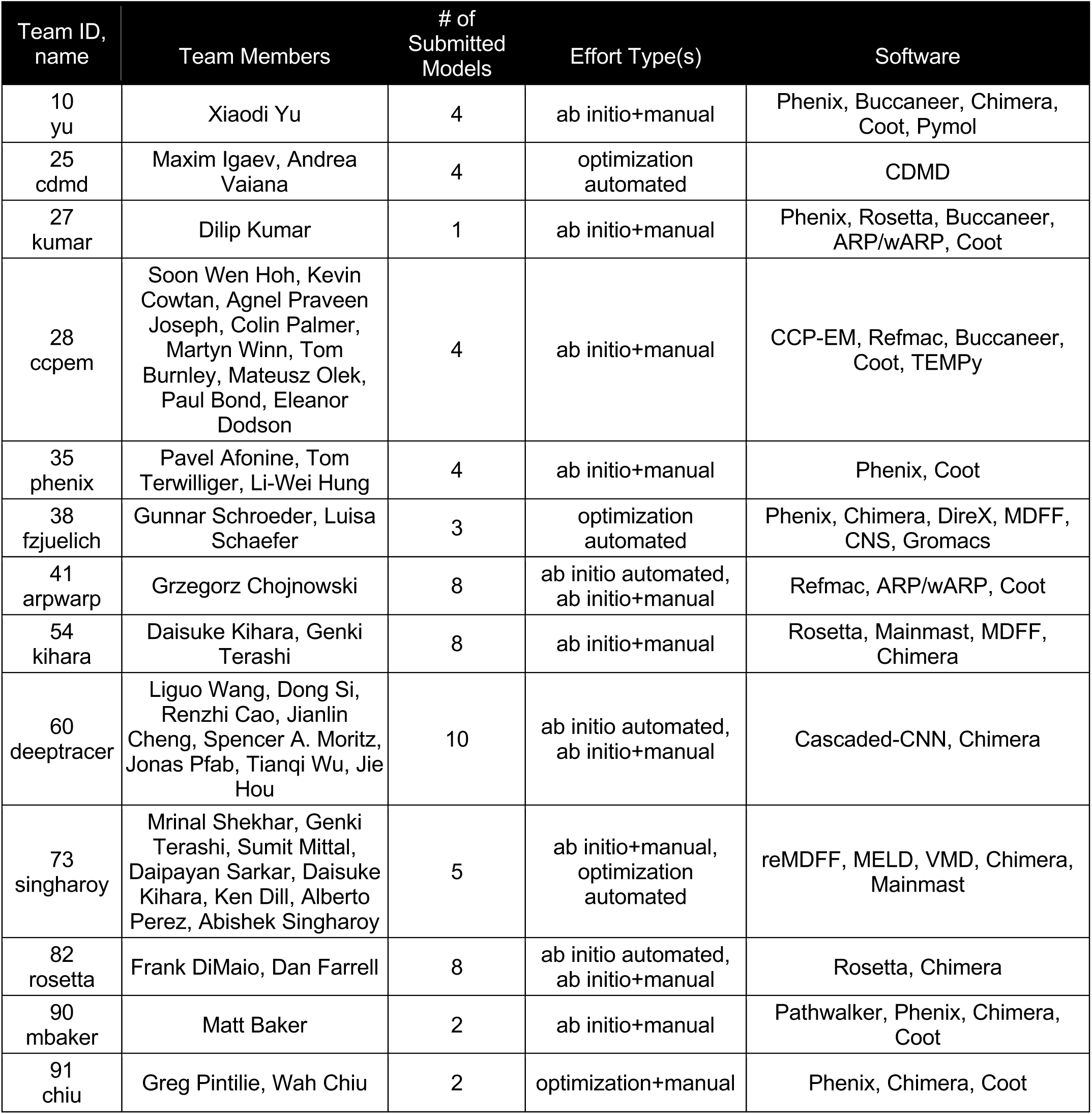
Participating Modeling Teams

Submitted models were evaluated as in the previous Challenge^12^ with multiple metrics in each of four tracks: Fit-to-Map, Coordinates-only, Comparison-to-Reference, and Comparison-among-Models (Figure 3c, Table II). The selected metrics include many already in common use, as well as several introduced via this Challenge.

**Table II.** Evaluated Metrics

|  | Metric Class | Package <u>Metric</u> Definition |
| --- | --- | --- |
| Fit-to-Map | Correlation Coefficient, all voxels | Phenix <u>CCbox</u> full grid map vs model-map density correlation coefficient<sup>18</sup><br>TEMPy <u>CCC</u> full grid map vs model-map density correlation coefficient<sup>23</sup> |
|  | Correlation Coefficient, selected voxels | Phenix <u>CCmask</u> map vs model-map density, only modelled regions<sup>18</sup><br>Phenix <u>CCpeaks</u> map vs model-map density, only high-density map and model regions<sup>18</sup><br>TEMPy <u>CCC_OV</u> map vs model-map density, overlapping map and model regions<sup>25</sup><br>TEMPy <u>SMOC</u> Segment Manders' Overlap, map vs model-map density, only modelled regions<sup>25</sup> |
|  | Correlation Coefficient, other density function | TEMPy <u>LAP</u> map vs model-map Laplacian filtered density (partial 2<sup>nd</sup> derivative)<sup>22</sup><br>TEMPy <u>MI</u> map vs model-map Mutual Information entropy-based function<sup>22</sup><br>TEMPy <u>MI_OV</u> map vs model-map Mutual Information, only modelled regions<sup>25</sup> |
|  | Correlation Coefficient, atom positions | Chimera/MAPQ <u>Qscore</u> map density at each modeled atom vs reference Gaussian density function<sup>14</sup> |
|  | Fourier Shell Correlation | Phenix <u>FSC05</u> Resolution (distance) of Map-Model FSC curve read at point FSC=0.5<sup>18</sup><br>CCPEM/Refmac <u>FSCavg</u> FSC curve area integrated to map resolution limit<sup>19,56</sup> |
|  | Atom Inclusion | EMDB/VisualAnalysis <u>AI_all</u> Atom Inclusion, percentage of all atoms inside depositor-provided density threshold<sup>20</sup><br>TEMPy <u>ENV</u> Atom Inclusion in envelope corresponding to sample MW; penalizes unmodeled regions<sup>22</sup> |
| | Side Chain Density | Phenix <u>EMRinger</u> evaluates backbone positioning by sampling map density around Cγ-atom ring-paths for non-branched residues<sup>21</sup> |
| Coordinates-only | Configuration | Phenix <u>Bond</u> Root-mean-square deviation (RMSD) of bond lengths<sup>27</sup><br>Phenix <u>Angle</u> RMSD of bond angles<sup>27</sup><br>Phenix <u>Chiral</u> RMSD of chiral centers<sup>27</sup><br>Phenix <u>Planar</u> RMSD of planar group planarity<sup>27</sup><br>Phenix <u>Dihedral</u> RMSD of dihedral angles<sup>27</sup> |
| | Clashes | MolProbity <u>Clashscore</u> Number of steric overlaps $\geq 0.4\text{\AA}$ per 1000 atoms<sup>26</sup> |
| | Conformation | MolProbity <u>Rotamer</u> sidechain conformation outliers<sup>26</sup><br>MolProbity <u>Rama</u> Ramachandran $\phi, \psi$ mainchain conformation outliers<sup>26</sup><br>MolProbity <u>CaBLAM_3D</u> outliers of CO-CO virtual dihedral incompatible with 2 surrounding Ca-Ca virtual dihedrals<sup>28</sup><br>MolProbity <u>Calpha</u> 3D outliers of 2 Ca virtual mainchain dihedrals plus Calpha virtual bond angle<sup>28</sup> |
| vs. Reference Model(s) | Atom Superposition | LGA <u>GDT-TS</u> Global Distance Test Total Score, average % of model Ca that superimpose with reference Ca, multiple distance cutoffs<sup>29</sup><br>LGA <u>GDC</u> Global Distance Calculation, average % of all model atoms that superimpose with reference, multiple distance cutoffs<sup>29</sup><br>OpenStructure/QS <u>CaRMSD</u> Root-Mean-Square Deviation of Ca atoms<sup>31</sup> |
|  | Interatomic Distances | LDDT <u>LDDT</u> Local Difference Distance Test, superposition-free comparison of all-atom distance maps between model and reference<sup>30</sup> |
|  | Contact Area | CAD <u>CAD</u> Contact Area Difference, superposition-free measure of differences in interatom contacts<sup>32</sup><br>HBPLUS<sup>57</sup> <u>HBPRC</u>>6, Hydrogen bond precision, nonlocal. fraction of correctly placed hydrogen bonds in residue pairs with > 6 separation in sequence. |
|  | Atom Superposition, Multiple | <u>DAVIS QA</u> average of pairwise LGA GDT-TS scores among submitted models<sup>33</sup> |

Metrics to evaluate global Fit-to-Map included Map-Model Fourier Shell Correlation (FSC)^17^ as encoded in Phenix^18^, Refmac FSC average^19^, EMDB atom inclusion^20^, EMRinger^21^, multiple Map vs. Model density-based correlation scores from TEMPy^22-25^, Phenix^18^, and the recently introduced Q-score to assess atom resolvability^14^.

Metrics to evaluate overall Coordinates-only quality included Clashscore, Rotamer outliers, and Ramachandran outliers from MolProbity^26^, as well as standard geometry measures (bond, bond angle, chirality, planarity, and dihedral angle RMSDs) from Phenix^27^. PDB currently uses each of these validation measures, based on community recommendations^6-8^. New in this round was MolProbity CaBLAM, which evaluates protein backbone conformation across multiple residues using novel virtual dihedral angle definitions^28^.

Metrics assessing the similarity of a model to a reference structure included Global Distance Test total score^29^, Local Difference Distance Test^30^, CaRMSD from OpenStructure/QS^31^, and Contact Area Difference^32^. Davis-QA was used to measure similarity among submitted models^33^. All of these measures are widely used in CASP competitions^33^.

Several metrics were also evaluated at the per-residue level: Fit-to-Map: EMRinger, Q-score, EMDB atom inclusion, TEMPy SMOC, and Phenix CCbox; Coordinates-only: Clashes, Ramachandran outliers, and CaBLAM.

Evaluated metrics are tabulated with brief definitions in Table II; extended descriptions are provided in Online Methods.

An evaluation system website with interactive tables, plots and tools (Figure 3d) was established in order to organize and enable analysis of the Challenge results and to make the results accessible to all participants (<u>model-compare.emdataresource.org</u>).

### Overall and local quality of models

The vast majority of submitted models scored well, landing in “acceptable” regions for metrics in each of the evaluation tracks, and in many cases performing better than the associated reference structure which served as a control (Supplementary Figure 1). For teams that submitted *ab initio* models, additional manual adjustment was beneficial, particularly for models built into the two lower resolution targets. In general, the best scoring models were produced by well-established methods and experienced modeling practitioners.

Evaluation exposed four fairly frequent issues: mis-assignment of peptide-bond geometry, misorientation of peptides, local sequence misalignment, and failure to model associated ligands. Sidechain model quality was not specifically assessed in this round.

Two-thirds of the submitted models had one or more peptide-bond geometry errors (Supplementary Figure 2).

At resolutions near 3 Å or in weak local density, the carbonyl O protrusion disappears into the tube of backbone density (Figure 2), and *trans* peptide bonds are more readily modeled in the wrong orientation. If ϕ,ψ values are explicitly refined, adjacent side chains can be pushed further in the wrong direction instead of fixing the underlying problem. Such cases are not flagged as Ramachandran outliers but they are still recognized by CaBLAM^34^.

Sequence misthreadings misplace specific chemical groups over very large distances. The misalignment can be recognized by local Fit-to-Map criteria, with ends flagged by CaBLAM, bad geometry, *cis*-nonPro peptides, and clashes (Supplementary Figure 3).

The ADH map contains tightly bound ligands: an NADH cofactor as well as two zinc ions per subunit, with one zinc in the active site and the other in a spatially separate site where the metal coordinates with four cysteine residues^15^. A number of models lacking these ligands had considerable local modeling errors, sometimes even mistracing the backbone (Supplementary Figure 4).

**Figure 4.**
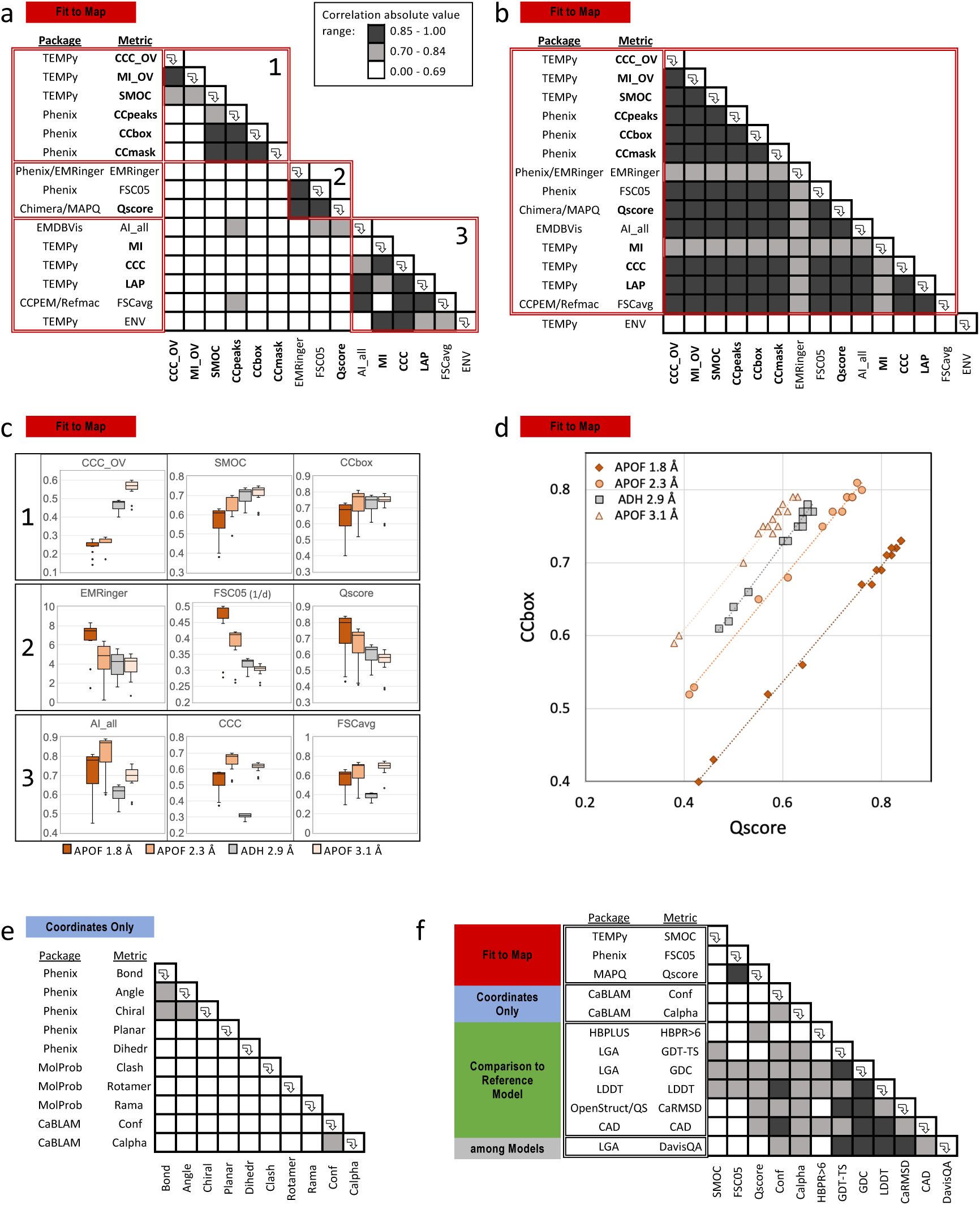
Evaluation of metrics. Model metrics (Table II) were compared with each other to assess how similarly they performed in scoring the Challenge models. (a-d) Fit-to-Map metrics were compared with each other to assess how similarly they performed in scoring the Challenge models. Their similarity was evaluated in two ways: (a) Pairwise correlation coefficients were calculated for all models across all map targets (n=63); (b) Average correlation per target was calculated (separate correlation coefficients for each map target were averaged). In both, pairs of metrics that are strongly similar in performance are indicated by dark shading (high correlation/black: 0.85-1.0; moderately high correlation/grey: 0.7-0.84). Those that perform very differently (poor correlation) are indicated with no shading. Correlation-based metrics are identified by bold labels. The ordering of metrics in (a) is based on hierarchical cluster analysis of all-model correlation values (see Methods). Three red-outlined boxes along the table diagonal correspond to identified clusters (#1-3); labels at left are also boxed in red according to these clusters. For ease of comparison, the ordering of metrics in (b) is identical to (a). The red-outlined box in (b) identifies the single cluster identified by hierarchical cluster analysis of the per-target correlation averages. This one cluster includes all metrics but ENV. In (c), representative score distributions for nine metrics, three from each cluster in (a), are plotted for each map target. These plots illustrate the systematic differences in scoring per map target that are responsible for the division of the evaluated metrics into the three clusters. In Cluster 1, score distributions are lowest for the highest resolution target and increase as map resolution decreases. Cluster 2 metrics have the opposite trend: score distributions are highest for the highest resolution target and decrease as map resolution decreases. Cluster 3 metrics have a mixed trend with respect to map resolution, but uniformly have lower score distributions for ADH map target models relative to all three APOF map target models. See the main text for discussion of the most likely factors behind these trends. In (d), scores for one representative pair of metrics that belong to different clusters in (a) but to the same cluster in (b) are plotted against each other. As highlighted by the diagonal lines representing linear fits to scores for map target, both metrics (CCbox from Cluster 1 and Qscore from Cluster 2) perform very similarly to each other within any one map target. Different sensitivities of the two metrics to map-specific factors give rise to the separate and parallel spacings of the four diagonal lines, with score ranges on different relative scales for each target. (e) Coordinates-only and (f) Comparison-to-Reference metrics comparisons. Analogously to (a) and (b), the Pearson correlation coefficient was determined for each metric pair for all submitted model scores (n=63) across all map targets. In (f), Comparison-to-reference metrics are contrasted with each other as well as with several Fit-to-Map and Coordinates-only metrics.

Although there was evidence for ordered water in the higher resolution APOF maps^14^, only two groups elected to model water oxygen atoms in their submissions. Model submissions were also split approximately 50:50 for the following practices: (1) inclusion of predicted hydrogen atom positions and (2) refinement of isotropic B-factors. Although near-atomic cryo-EM maps do not have a sufficient level of detail to directly identify hydrogen atom positions, inclusion of predicted H-atom positions can be useful for identifying model steric properties such as H-bonds or clashes^26^. Where provided, refined B-factors modestly improved Fit-to-Map scores against the highest resolution map target (APOF 1.8 Å) but had little to no benefit against lower resolution map targets.

### Evaluating Metrics: Fit-to-Map

Fit-to-Map metrics (Table II, red section) were systematically compared using score distributions of the submitted models (Figure 4a-d). For APOF targets, subunit models were evaluated against masked subunit maps, whereas for the ADH target, dimeric models were evaluated against the full sharpened cryo-EM map (Figure 2d). To control for the impact of hydrogen atom inclusion or isotropic B-factor refinement on different subsets of Fit-to-Map metrics, all evaluated scores were produced with hydrogen atoms removed and with B-factors set to zero.

Score distributions were first evaluated *for all 63 models across all four Challenge targets*. Unexpectedly, a wide diversity in performance was observed, with poor correlations between most pairs of metrics (Figure 4a). This means that a model that scored well relative to all 62 others using one metric may have a much poorer ranking using another metric. A hierarchical cluster analysis identified three distinct clusters of similarly performing metrics (Figure 4a, boxes 1-3).

The observed sparse correlations and clustering of the Fit-to-Map metrics can be readily understood by considering their per target score distribution ranges, which differ substantially from each other (Figure 4c). The three clusters identify sets of metrics that share similar trends (Fig. 4c, panels 1-3).

#### Cluster 1

metrics (Figure 4c, panel 1) share the trend of decreasing score values with increasing map target resolution. The cluster consists of six correlation measures, three from TEMPy^22-25^ and three from Phenix^18^. Each evaluates a model’s fit to the map in a similar way: by correlating a calculated model-map density with the experimental map density. In most cases (5 of 6), correlation is performed following model-based masking of the experimental map. The observed trend arises at least in part because as map resolution increases, the level of detail that a model-map must faithfully replicate in order to achieve a high correlation score must also increase.

#### Cluster 2

metrics (Figure 4c, panel 2) share the inverse trend: score values improve with increasing map target resolution. Cluster 2 metrics consist of Phenix Map-Model FSC=0.5^18^, Qscore^14^, and EMRinger^21^. The observed trend is expected: by definition each metric assesses a model’s fit to the experimental map in a manner that is sensitive to map resolution.

#### Cluster 3

metrics (Figure 4c, panel 3) share a different trend: score values are significantly lower for ADH relative to APOF map targets. These measures include three unmasked correlation functions from TEMPy^22-25^, Refmac FSCavg^19^, EMDB Atom Inclusion^20^ and TEMPy ENV^22^. All of these measures consider the full experimental map without masking, so can therefore be sensitive to background noise. Background noise was substantial in the unmasked ADH map and minimal in the masked APOF maps (Figure 2d).

Score distributions were also evaluated for how similarly they performed *per target*, and in this case most metrics were strongly correlated with each other (Figure 4b). This means that within any single target, a model that scored well relative to all others using one metric also fared well using nearly every other metric. This situation is illustrated by comparing scores for two different metrics, CCbox from Cluster 1 and Q-score from Cluster 2 (Figure 4d). The plot’s four diagonal lines demonstrate that the scores are tightly correlated with each other within each map target. But as described above in the analyses of Clusters 1 and 2, the two metrics each have different sensitivities to map-specific factors. It is these different sensitivities that give rise to the separate and parallel spacings of the four diagonal lines, indicating score ranges on different relative scales for each target.

One Fit-to-Map metric showed poor correlation with all others in the per target analysis: TEMPy ENV (Figure 4b). ENV scores were poorly distributed with most models very close to the maximum possible value (1.0). ENV evaluates atom positions relative to a density threshold that is determined from the sample molecular weight. At near-atomic resolution this threshold is overly generous and tends to include all modelled atoms. TEMPy Mutual Information (MI) and EMRinger also diverged somewhat from the other metrics (Figure 4b). Within each target, all MI scores were essentially identical to each other. This behavior may reflect a strong influence of background noise, since MI_OV, MI’s masked version, yielded distributed scores that correlated well with other measures. As noted previously^21^, EMRinger follows similar trends with other measures but yields distinct distributions owing to its focus on backbone placement.

Collectively these results reveal that multiple factors such as experimental map resolution, presence of background noise, and density threshold selection can strongly impact Fit-to-Map score values, depending on the chosen metric.

### Evaluating metrics: Coordinates-only and vs-Reference

Metrics to assess model quality based on Coordinates-only (Table II, blue section), as well as Comparison-to-Reference and Comparison-among-Models (Table II, green and grey sections) were also evaluated and compared (Figure 4e-f).

Most of the Coordinates-only metrics were poorly correlated with each other (Figure 4e), with the exception of bond, bond angle, and chirality RMSD, which form a small cluster. Interestingly, Ramachandran outlier score, which is widely used to assess protein backbone conformation, was poorly correlated with all other Coordinate-only measures, including the novel CaBLAM scores^28^. Score distributions explain this in part: more than half (33) of submitted models had zero Ramachandran outliers, while only four had zero CaBLAM Conformation outliers (we note that Ramachandran statistics are increasingly used as restraints^35,36^). These results support the concept of CaBLAM as a new informative score for validating backbone conformation^28^.

The CaBLAM Conformation and C-alpha measures, while orthogonal to other Coordinate-only measures, were unexpectedly found to perform very similarly to Comparison-to-Reference metrics; several Fit-to-Map metrics also performed somewhat similarly to Comparison-to-Reference metrics (Figure 4f). The similarity likely arises because the worst modeling errors in this Challenge were sequence and backbone conformation mis-assignments. These errors were equally flagged by CaBLAM, which compares models against statistics of high-quality structures from the PDB, and the Comparison-to-Reference metrics, which compare models directly against a high-quality reference. To a somewhat lesser extent these modeling errors were also flagged by Fit-to-Map metrics.

### Evaluating metrics: local scoring

As part of the evaluation pipeline, residue-level scores were calculated in addition to overall scores. Five Fit-to-Map metrics either considered masked density for both map and model around the evaluated residue (Phenix CCbox^18^, TEMPy SMOC^24^), density profiles at non-hydrogen atom positions (Qscore^14^), density profiles of non-branched residue Cγ-atom ringpaths (EMRinger^21^), or density values at non-hydrogen atom positions relative to a chosen threshold (EMDB Atom Inclusion^20^). In two of the five, residue-level scores were obtained as sliding-window averages over multiple contiguous residues (SMOC: 9-residues; EMRinger: 21-residues).

Residue-level correlation analyses similar to those described above showed that local fit-to-map scores diverged more than their corresponding global scores. Residue-level scoring was most similar across the evaluated metrics for high resolution maps. This observation suggests that the choice of method for scoring residue-level fit becomes less critical at higher resolution, where maps tend to have stronger density/contrast around atom positions.

A case study of a local modeling error in one of the APOF 2.3 Å models (Supplementary Figure 3) showed that EMDB Atom Inclusion^20^, Phenix CCbox^18^, and Qscore^14^ measures produced significantly lower (worse) scores within a 4-residue α-helical misthread relative to correctly assigned flanking residues. In contrast, the two sliding-window-based metrics were largely insensitive (a more recent version of TEMPy offers single residue analysis (SMOCd) and adjustable window analysis based on map resolution (SMOCf)^37^). At near-atomic resolution, single residue fit-to-map evaluation methods are likely to be more useful than windowing methods for identifying local modelling issues.

Residue-level Coordinate-only metrics (Supplementary Figure 3), Comparison-to-Reference metrics and Comparison-among-Models metrics (not shown) were also evaluated for the same modeling error. The MolProbity server^26,28^ flagged the problematic 4-residue misthread via CaBLAM, cis-Peptide, clashscore, bond, and angle scores, but all Ramachandran scores were either favored or allowed. The Comparison-to-Reference LDDT and LGA local scores and the Davis-QA model consensus score also strongly flagged this error. The example demonstrates the value of combining multiple orthogonal measures to identify geometry issues, and further highlights the value of CaBLAM as a novel, orthogonal measure for validation of backbone conformation.

### Group performance

Group performance was examined by modeling category and target by combining Z-scores from metrics determined to be meaningful in the analyses described above (see Methods and Supplementary Figure 5).

For *ab initio* modeling, lower resolution targets were more challenging for some groups. For the higher resolution APOF 1.8 Å and 2.3 Å targets, six groups (10, 28, 35, 41, 73, 82, see Table I ids) did very well (Z ≥ 0.3), and a seventh (54, models 2) was a runner-up. For the lower resolution APOF 3.1 Å and ADH 2.9 Å targets, a slightly different six groups (10, 27, 28, 35, 73, did very well and another two (41, 90) were runners-up. A wide variety of map density features and algorithms to produce a model, and most were quite successful yet allowing a few mistakes, often in different places (see Supplementary Figures 2-4). For practitioners, it might be beneficial to compare/combine models from several *ab initio* methods to come up with a better initial model for subsequent refinement. Note that the performance results are specific to the Challenge task and may not be directly applicable to other modeling scenarios.

As for optimization-based modeling, all made improvements, but sample size was too small to produce a rating.

## Discussion

This 3rd Model Challenge round has demonstrated that cryo-EM maps with resolution ≤ 3 Å and from samples with limited conformational flexibility, have excellent information content, and automated methods are able to generate fairly complete models from such maps, needing only small amounts of manual intervention to be finalized (but some is always needed). Modeling could readily be accomplished within a month, the time-period of this challenge. This outcome represents a great advance over the previous challenges.

Inclusion of three maps in a resolution series enabled controlled evaluation of metrics by resolution. Inclusion of a completely different map as the fourth target provided a useful additional control. These target selections enabled observation of important trends that otherwise could have been missed. In a recent evaluation of predicted models against several ∼3 Å cryo-EM maps in the CASP13 competition, TEMPy and Phenix Fit-to-Map correlation measures performed very similarly^37^. In this Challenge, because the chosen map targets covered a wider resolution range and had more variability in background noise, the same measures were found to have distinctive, map target feature-sensitive performance profiles.

The majority of submitted models were overall either equivalent to or better than the reference model in terms of the fit of their atomic coordinates to the target map. This achievement reflects significant advances in the development of modeling tools relative to the state presented a decade ago in our first Model Challenge^9^. However, several factors beyond atom positions that become important for accurate modelling at near-atomic resolution were not uniformly addressed: only half of the submitted models included refinement of atomic displacement factors (B-factors), and a minority of modellers attempted to fit water or bound ligands.

Fit-to-Map measures were found to be sensitive to different physical properties of the map, including experimental map resolution and background noise level, as well as input parameters such as density threshold. Coordinates-only measures were found to be largely orthogonal to each other, while Comparison-to-Reference measures were generally well correlated with each other.

The cryo-EM modeling community as represented by the Challenge participants have introduced a number of metrics to evaluate cryo-EM models with sound biophysical basis. We find that some of them are correlated to each other and to the resolution of the map, while others are not. Based on our careful analyses of these metrics and their relationships, we make four recommendations regarding validation practices for cryo-EM models of proteins determined at near-atomic resolution as studied here between 3.1 Å and 1.8 Å, a rising trend for cryo-EM (Figure 1).

### Recommendation 1

For researchers optimizing a model against a single map, nearly any of the evaluated global fit-to-map metrics (Table II) can be used to evaluate progress because they are all largely equivalent in performance. Exception: TEMPy ENV is more appropriate for medium to low resolution (>4 Å).

### Recommendation 2

To flag issues with local (per residue) fit to a map, metrics that evaluate single residues such as CCbox, Qscore, and EMDB Atom Inclusion are more suitable than those using sliding window averages over multiple residues.

### Recommendation 3

The ideal Fit-to-Map metric for *archive-wide ranking* will be insensitive to map background noise (appropriate masking or alternative data processing can help), will not require input of estimated parameters that affect score value (*e*.*g*., resolution limit, threshold). and will yield overall better scores for maps with trustworthy higher-resolution features. The three Cluster 2 metrics identified in this Challenge (Figure 4a) meet these criteria.

- Map-Model FSC^17,18^ is already in common use ^13^, and can be compared with the experimental map’s independent half-map FSC curve.
- Global EMRinger score^21^ can assess non-branched protein side chains.
- Q-scoreis a relatively new correlation metric that can be used both globally and locally for validating non-hydrogen-atom x,y,z positions.^14^.

Other Fit-to-map metrics may be rendered suitable for archive-wide comparisons through conversion of raw scores to Z-scores over narrow resolution bins, as is currently done by the PDB for some X-ray-based metrics^7,38^.

### Recommendation 4

CaBLAM statistical measures and MolProbity cis-peptide detection^28^ are useful to detect protein backbone conformation issues. These are valuable new tools for cryo-EM protein structure validation, particularly since maps at typical resolutions (2.5 - 4.0 Å; Figure 1) may not resolve backbone carbonyl oxygens (Figure 2).

The 2019 Model “Metrics” Challenge was more successful than previous challenges because more time could be devoted to analysis and because infrastructure for model collection, processing and assessment is now established. EMDR plans to sponsor additional model challenges in order to continue promoting development and testing of cryo-EM modeling and validation methods. Future challenge topics are likely to cover medium resolution (3 to 4 Å), particle heterogeneity, membrane proteins, ligand modeling, nucleic acids, and models derived from tomograms.

## Online Methods

### Challenge Process and Organization

Informed by previous Challenges^9,10,12^, the 2019 Model Challenge process was significantly streamlined in this round. In March, a panel of advisors with expertise in cryo-EM methods, modeling, and/or model assessment was recruited. The panel worked with EMDR team members to develop the challenge guidelines, identify suitable map targets from EMDB and reference models from PDB, and recommend the metrics to be calculated for each submitted model.

The Challenge rules and guidance were as follows: (1) *Ab initio* modeling is encouraged but not required. For optimization studies, any publicly available coordinate set can be used as the starting model. (2) Regardless of the modeling method used, submitted models should be as complete and as accurate as possible (*i*.*e*., equivalent to publication-ready). (3) For each target, a separate modeling process should be used. (4) Fitting to either the unsharpened/unmasked map or one of the half-maps is strongly encouraged. (5) Submission in mmCIF format is strongly encouraged.

Members of cryo-EM and modeling communities were invited to participate in mid-April 2019; details were posted on the challenges website (<u>challenges.emdataresource.org</u>). Models were submitted by participant teams between May 1 and May 28, 2019. For apoferritin (APOF) targets, coordinate models were submitted as single subunits at the position of a provided segmented density consisting of a single subunit. Alcohol dehydrogenase (ADH) models were submitted as dimers. For each submitted model, metadata describing the full modeling workflow were collected via a Drupal webform, and coordinates were uploaded and converted to PDBx/mmCIF format using PDBextract^39^. Model coordinates were then processed for atom/residue ordering and nomenclature consistency using PDB annotation software (Feng Z., https://swtools.rcsb.org/apps/MAXIT) and additionally checked for sequence consistency and correct position relative to the designated target map. Models were then evaluated as described below (Model Evaluation System).

In early June, models, workflows, and initial calculated scores were made available to all participants for evaluation, blinded to modeler team identity and software used. A 2.5-day workshop was held in mid-June at Stanford/SLAC to review the results, with panel members attending in person. All modeling participants were invited to attend remotely and present overviews of their modeling processes and/or assessment strategies. Recommendations were made for additional evaluations of the submitted models as well as for future challenges. Modeler teams and software were unblinded at the end of the workshop. In September, a virtual follow-up meeting with all participants provided an overview of the final evaluation system after implementation of recommended updates.

### Modeling Software

Modelling teams created *ab initio* models or optimized previously known models available from the PDB. *Ab initio* software included ARP/wARP^40^, Buccaneer^41,42^, Cascaded-CNN^43^, Mainmast^44, Terashi 2020^, Pathwalker^45^, and Rosetta^46^. Optimization software included CDMD^47^, CNS^48^, DireX^49^, Phenix^27^, REFMAC^19^, MELD^50^, MDFF^51^, and reMDFF^52^. Participants made use of VMD^53^, Chimera^54^, and COOT^35^ for visual evaluation and/or manual model improvement of map-model fit. See Table I for software used by each modeling team.

### Model Evaluation System

The evaluation system for 2019 Challenge (<u>model-compare.emdataresource.org</u>) was built on the basis of the 2016/2017 Model Challenge system^12^, updated with several new evaluation measures and analysis tools. Submitted models were evaluated for >70 individual metrics in four tracks: Fit-to-Map, Coordinates-only, Comparison-to-Reference, and Comparison-among-Models. A detailed description of the updated infrastructure and each calculated metric is provided as a help document on the model evaluation system website.

For brevity, a representative subset of metrics from the evaluation website are discussed in this paper. The selected metrics are listed in Table II, and are further described below. All scores were calculated according to package instructions using default parameters.

#### Fit-to-Map

The evaluated metrics included several ways to measure the correlation between map and model density^55^, as implemented in TEMPy^22-25^ v.1.1 (CCC, CCC_OV, SMOC, LAP, MI, MI_OV) and the Phenix^27^ v.1.15.2 map_model_cc module^18^ (CCbox, CCpeaks, CCmask). These methods compare the experimental map with a model map produced on the same voxel grid, integrated either over the full map or over selected masked regions. The model-derived map is generated to a specified resolution limit by inverting Fourier terms calculated from coordinates, B-factors, and atomic scattering factors. Some measures compare density-derived functions instead of density (Mutual Information, Laplacian^22^).

The newly introduced Q-score (MAPQ v1.2^14^ plugin for UCSF Chimera^54^ v.1.11) uses a real-space correlation approach to assess the resolvability of each model atom in the map. Experimental map density is compared to a Gaussian placed at each atom position, omitting regions that overlap with other atoms. The score is calibrated by the reference Gaussian, which is formulated so that a highest score of 1 would be given to a well-resolved atom in a map at ∼1.5 Å resolution. Lower scores (down to -1) are given to atoms as their resolvability and the resolution of the map decreases. The overall Q-score is the average value for all model atoms.

Measures based on Map-Model FSC curve, atom inclusion, and protein side chain rotamers were also compared. Phenix Map-Model FSC is calculated using a soft mask and is evaluated at FSC=0.5^18^. REFMAC FSCavg^19^ (module of CCPEM^56^ v1.4.1) integrates the area under the Map-Model FSC curve to a specified resolution limit^19^. EMDB Atom Inclusion determines the percentage of atoms inside the map at a specified density threshold^20^. TEMPy ENV is also threshold-based and penalizes unmodeled regions^22^. EMRinger (module of Phenix) evaluates backbone positioning by measuring the peak positions of unbranched protein C_γ_ atom positions versus map density in ring-paths around C_α_-C_β_ bonds^21^.

#### Coordinates-only

Standard measures assessed local configuration (bonds, bond angles, chirality, planarity, dihedral angles; Phenix model statistics module), protein backbone (MolProbity Ramachandran outliers^26^; Phenix molprobity module) and side-chain conformations, and clashes (MolProbity rotamers outliers and clashscore^26^; Phenix molprobity module).

New in this Challenge round is CaBLAM^28^ (part of MolProbity and as Phenix cablam module), which employs two novel procedures to evaluate protein backbone conformation. In both cases, virtual dihedral pairs are evaluated for each protein residue *i* using C_α_ positions *i*-2 through *i*+2. To define CaBLAM outliers, the third virtual dihedral is between the CO groups flanking residue *i*. To define Calpha-geometry outliers, the third parameter is the Calpha virtual angle at *i*. The residue is then scored according to virtual triplet frequency in a large set of high-quality models from PDB^28^.

#### Comparison-to-Reference and Comparison-among-Models

Assessing the similarity of the model to a reference structure and similarity among submitted models, we used metrics based on atom superposition (LGA GDT-TS and GDC scores^29^ v.04.2019), interatomic distances (LDDT score^30^ v.1.2), and contact area differences (CAD^32^ v.1646). HBPLUS^57^ was used to calculate nonlocal hydrogen bond precision, defined as the fraction of correctly placed hydrogen bonds in residue pairs with > 6 separation in sequence.

DAVIS-QA determines for each model the average of pairwise GDT-TS scores among all other models^33^.

#### Local (per residue) Scores

Residue-level visualization tools for comparing the submitted models were also provided for the following metrics. Fit-to-Map: Phenix CCbox, TEMPy SMOC, Qscore, EMRinger, EMDB Atom Inclusion; Comparison-to-Reference: LGA and LDDT; Comparison-among-Models: DAVIS-QA.

### Metric Score Pairwise Correlations and Distributions

For pairwise comparisons of metrics, Pearson correlation coefficients (P) were calculated for all model scores and targets (N=63). For average per-target pairwise comparisons of metrics, P values were determined for each target and then averaged. Metrics were clustered according to the similarity score (1-|P|) using a hierarchical algorithm with complete linkage. At the beginning, each metric was placed into a cluster of its own. Clusters were then sequentially combined into larger clusters, with the optimal number of clusters determined by manual inspection. In the fit-to-map evaluation track, the procedure was stopped after three divergent score clusters were formed for the all-model correlation data (Figure 4a), and after two divergent clusters were formed for the average per-target clustering (Figure 4b).

Score distributions are represented in box-and-whisker format in Figure 4c. Each box represents the interquartile range (IQR) and is drawn between Q1 (25th percentile) and Q3 (75th percentile) values. The inner horizontal line represents the median value (excluding outliers). Whisker lines extend out to the highest and lowest measured scores that are within 1.5*IQR of each box end. Scores falling outside the 1.5*IQR limits are considered outliers and are separately plotted as dots.

### Controlling for Model Systematic Differences

As initially calculated, some Fit-to-Map scores had unexpected distributions, owing to differences in modeling practices among participating teams. For models submitted with all atom occupancies set to zero, occupancies were reset to one and rescored. In addition, model submissions were split approximately 50:50 for each of the following practices: (1) inclusion of hydrogen atom positions and (2) inclusion of refined B-factors. For affected fit-to-map metrics, modified scores were produced excluding hydrogen atoms and/or setting B-factors to zero. Both original and modified scores are provided in the web interface. Only modified scores were used in the pairwise metric comparisons described here.

### Evaluation of Group Performance

Rating of group performance was done using the Model Compare Pipeline/Comparative Analyses/Model Ranks (per target) tool on the Challenge evaluation website. The tool permits users, for a specified target and for all or a subcategory of models (e.g., *ab initio*), to calculate Z-scores for each individual model, using any combination of 47 of the evaluated metrics with any desired relative weightings. The Z-scores for each metric are calculated from all submitted models for that target. The metrics (weights) used to generate individual-model Z-scores were as follows:

#### Coordinates-only

CaBLAM outliers (0.5), Calpha-geometry outliers (0.3), and Clashscore (0.2). CaBLAM outliers and Calpha-geometry outliers had the best correlation with match-to-target parameters (Figure 5b), and clashscore is an orthogonal measure. Ramachandran and rotamer criteria were excluded since they are often restrained in refinement and are zero for many models.

#### Fit-to-Map

EMringer (0.3) and Q-score (0.3), Atom Inclusion-backbone (0.2), and SMOC (0.2). EMringer and Q-score were among the most promising model-to-map metrics, and the other two provide distinct measures.

#### Comparison-to-Reference

LDDT (0.9) and GDC_all (0.9) and HBPR>6 (0.2). LDDT is superposition-independent and local, while GDC_all requires superposition; H-bonding is distinct. Metrics in this category are weighted higher, because although the target models are not perfect, they are a reasonable estimate of the right answer.

Individual Z-scores for each model were then averaged across each group’s models on a given target, and further averaged across T1+T2 and across T3+T4, yielding overall Z-scores for high and low resolutions. The scores consistently showed 3 quite separate clusters: a good cluster at Z>0.3, an unacceptable cluster at Z<-0.3, and a small cluster near Z=0 (see Supplementary Figure 5). Other choices of metrics were tried, with very little effect on clustering.

Group 54 models were rated separately because they used different methods, their 2nd model versions were much better. Group 73’s second model on target T4 was not rated because the metrics are not set up to meaningfully evaluate an ensemble.

### Molecular Graphics

Molecular graphics images were generated using UCSF Chimera^54^ (Figure 2 and Supplementary Figure : maps with superimposed models) and KiNG^58^ (Supplementary Figures 2 and 4: maps with superimposed models and validation flags).

## Acknowledgements

EMDataResource is supported by the US National Institutes of Health (NIH)/National Institute of General Medical Science, R01GM079429.

The following additional grants are acknowledged for participant support. JSR and CW: NIH/ R35GM131883, NIH/P01GM063210.

AS: National Science Foundation (NSF)/MCB-1942763 (CAREER), NIH/R01GM095583. The Singharoy team used supercomputing resources of the OLCF at the Oak Ridge National Laboratory, which is supported by the Office of Science at DOE under Contract No. DE-AC05-00OR22725.

DKihara: NIH/ R01GM123055, NSF/DMS1614777, NSF/CMMI1825941, NSF/MCB1925643, Purdue Institute of Drug Discovery/DBI2003635.

JSF: NIH/R01GM123159.

MI: Max Planck Society German Research Foundation/IG 109/1-1.

ACV: Max Planck Society German Research Foundation /FOR-1805.

DKumar: NIH/R37AI36040 and Welch Foundation/Q1279 (PI: BVV Prasad).

DS: NSF/DBI2030381.

TB, CMP, MW: Medical Research Council MR/N009614/1.

APJ, MW: Wellcome Trust 208398/Z/17/Z.

KC: Biotechnology and Biological Sciences Research Council / BB/P000517/1.

MW: Biotechnology and Biological Sciences Research Council BB/P000975/1.

## Author Contributions

PDA, PVA, JF, FDM, JSR, PBR, HMB, WC, AK, CLL, GDP, MFS: expert panel that selected targets, reference models and assessment metrics, set challenge rules; attended face-to-face results review workshop. KZ generated the APF maps for the challenge. MAH provided the published ADH map. CLL: designed and implemented the challenge model submission pipeline, drafted initial manuscript. Authors as listed in Table I: Built and submitted models; presented modeling strategies at review workshop. AK: designed and implemented evaluation pipeline and website, calculated scores. AK, CLL, BM, MAH, JSR, CJW, PVA, JF: analyzed models and model scores. AP, ZW, MT, ADJ, GDP, PVA, CJW: contributed software, advice on use and scores interpretation. CLL, AK, GDP, JSR: drafted figures. AK, HMB, GDP, WC, MFS, MAH, JSR: contributed to manuscript writing. All authors: reviewed and approved final manuscript.

## Competing Interests

The authors declare no competing interests.

## Notes

https://model-compare.emdataresource.org/

https://challenges.emdataresource.org/

### Competing Interest Statement

The authors have declared no competing interest.

