## Supplementary material for "Outcomes of the 2019 EMDataResource model challenge: validation of cryo-EM models at near-atomic resolution": Supplmentary Figures 1-5

### Supplementary Figure 1. Model Score Distributions

Score distributions for all 63 models submitted to the challenge are shown on the following three pages (A: Fit-to-Map; B: Coordinates-only; C: Comparison-to-Reference). Parallel lanes display score distributions for each target and each evaluated metric. Scores are plotted horizontally as semi-transparent diamonds against a color-coded background to indicate worse (left, orange) and better (right, green) values. Where multiple scores overlap, the semi-transparent diamonds appear as a single darker, more opaque diamond. Reference model scores are indicated as red triangles.

A. Fit-to-Map

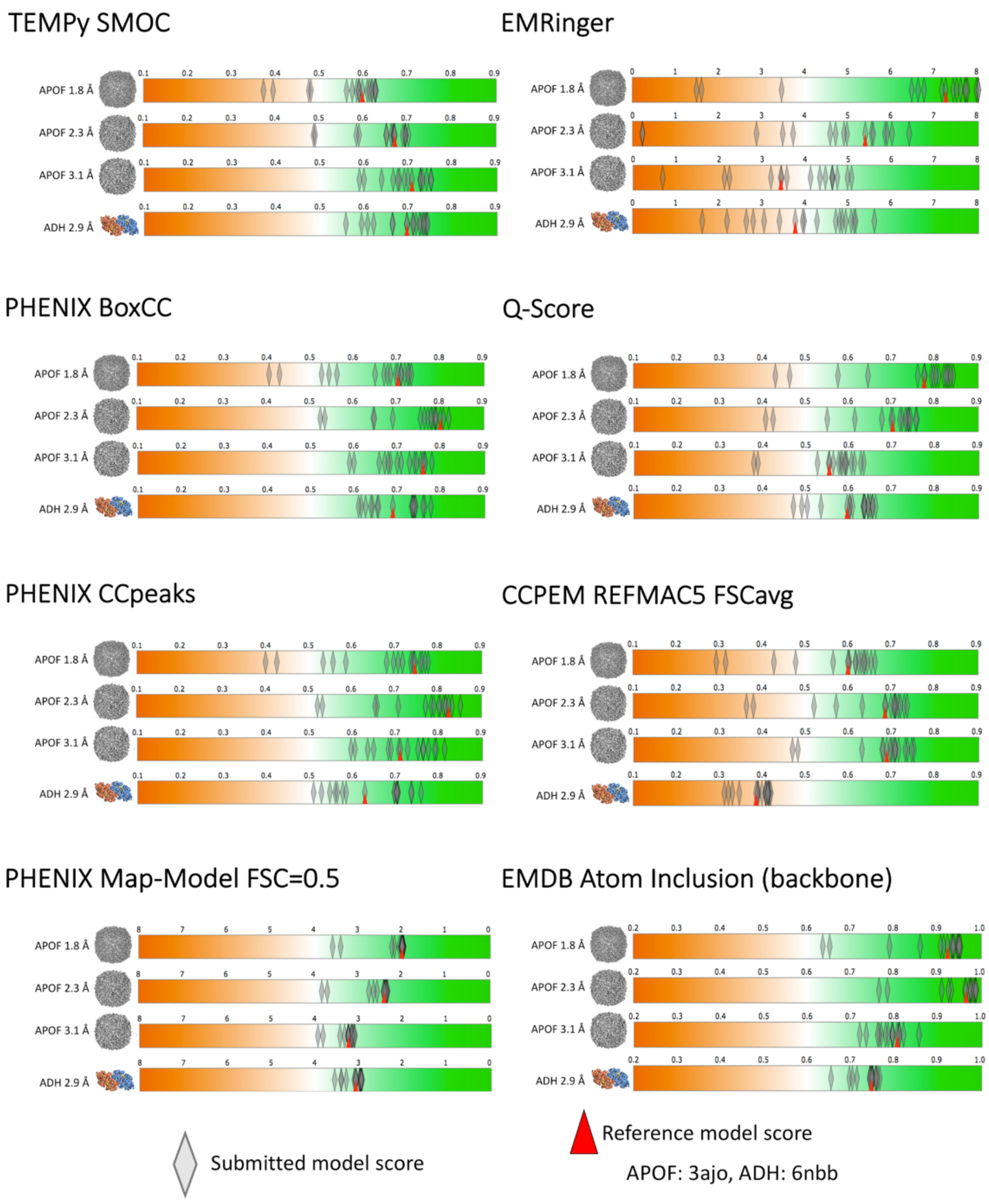
TEMPy SMOC
EMRinger
PHENIX BoxCC
Q-Score
PHENIX CCpeaks
CCPEM REFMAC5 FSCavg
PHENIX Map-Model FSC=0.5
EMDB Atom Inclusion (backbone)
APOF 1.8 Å
APOF 2.3 Å
APOF 3.1 Å
ADH 2.9 Å
Submitted model score
Reference model score
APOF: 3ajo, ADH: 6nbb

B. Coordinates-only

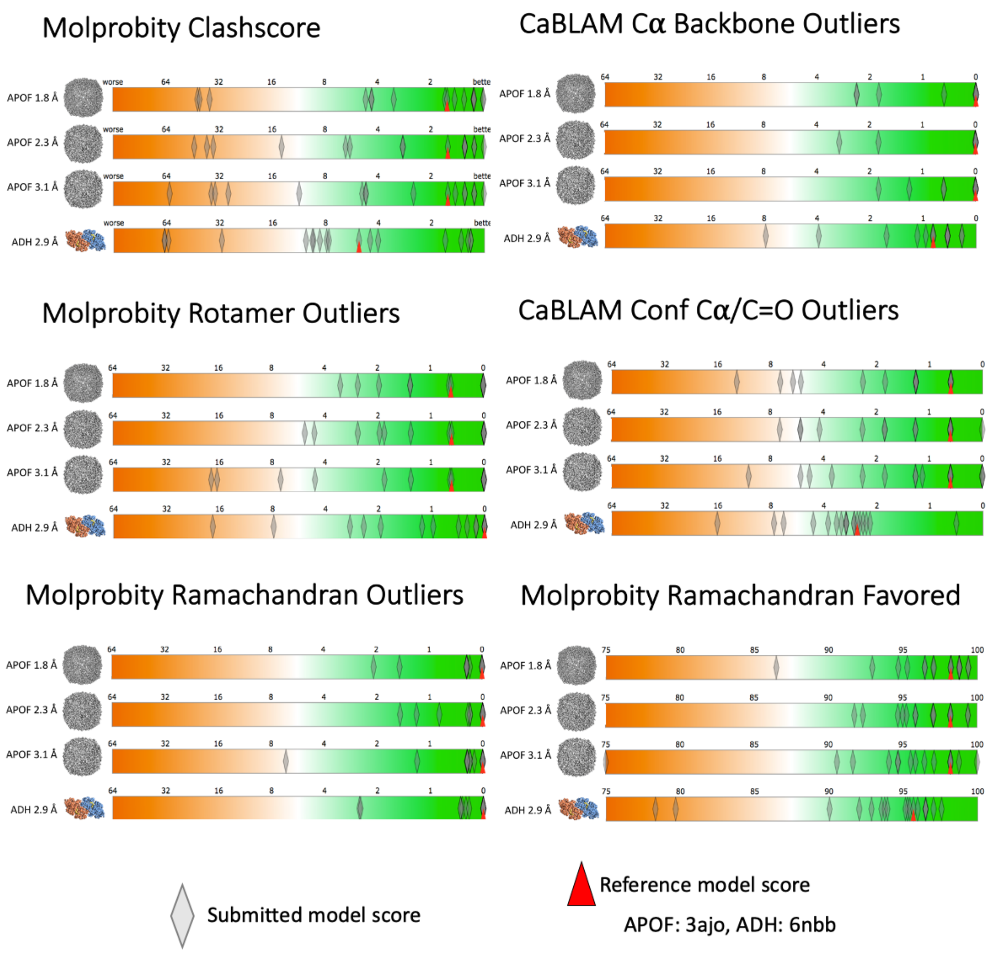

C. Comparison with Reference (monomeric)

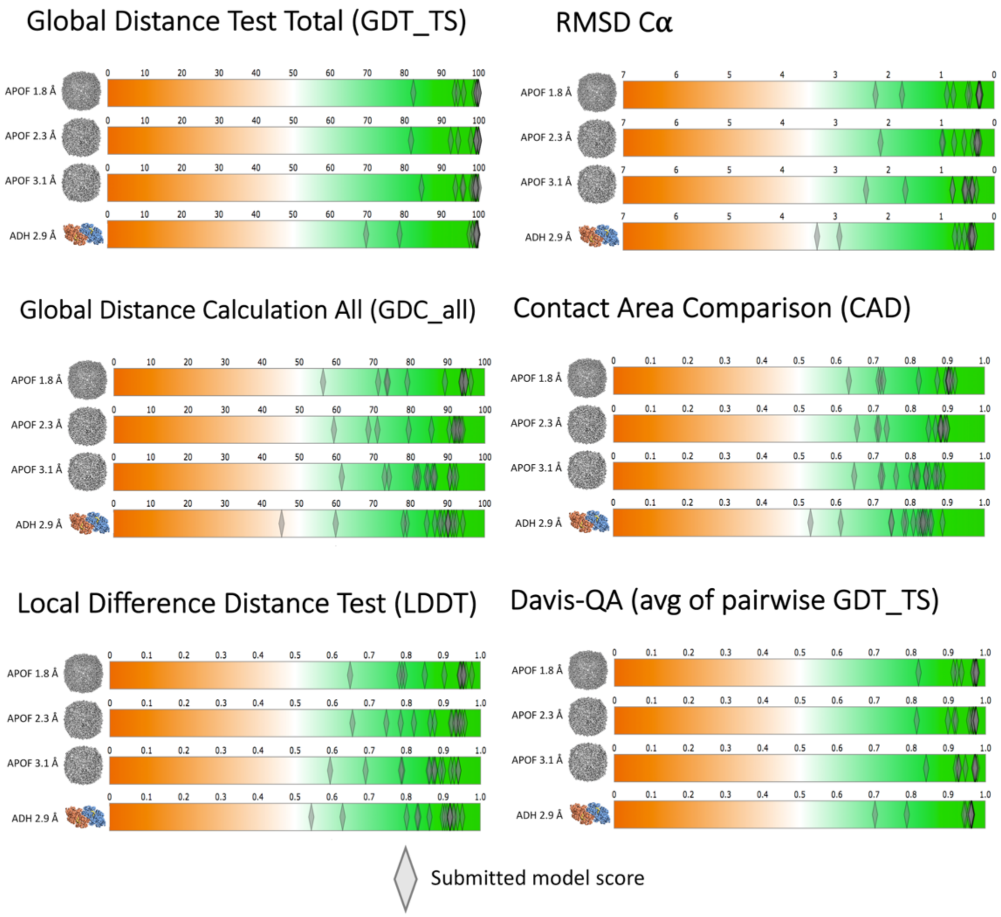

### Supplementary Figure 2. Evaluation of peptide bond geometry

All 63 Challenge models were evaluated using MolProbity[1,2]. The APOF and ADH reference models each have one *cis* peptide bond per subunit before a proline residue (cis-Pro). (a) Counts of peptides with each of the following conformational properties: cisP: cis peptide bond before proline, twistP: non-planar peptide bond (by >30°) before proline, cis-nonP: cis peptide bond before a non-proline, twist-nonP: non-planar peptide bond before a non-proline. Incorrect *cis*-nonPro instances usually occurred where the model was misfit (see Supplementary Figure 3), while incorrect *cis* or *trans* Pro instances usually produced extremely bad geometry. Values inconsistent with reference model scores are highlighted. Statistically, 1 in 20 proline residues are genuinely *cis*-Pro; only 1 in 3000 non-proline residues are genuinely *cis* (*cis*-nonPro), and strongly non-planar peptide bonds (>30°) are almost never genuine[29]. Model scores are organized by the submitting group (Gp #_# indicates group id as defined in Table I plus model #), and Target (T1-T4). Optimized models are shaded blue. Dashes indicate that no model was submitted for the specified target. Boxes with notations b-d indicate models illustrated in panels b-d. Only models 28_1 and 35_1 had all peptides correct for all 4 targets. (b) Correct cis peptide geometry for Pro A62 in two ADH (T4) models. (c) Incorrect trans peptide geometry, with huge clashes up to 1.25Å overlap (clusters of hot pink spikes), 2 CaBLAM outliers (magenta CO dihedral lines), and poor density fit. (d) Incorrect trans peptide geometry, with huge 1.9 Å Cβ deviation at Leu 61 (magenta ball) because of incorrect hand of Cα, and 2 CaBLAM outliers. Molecular graphics were generated using KiNG[3].

a

Model: cisP, twistP, cis-nonP, twist-nonP    not *ab initio*

| | | | | | |
|---|---|---|---|---|---|
| Gp 10_1 | T1: 1, 0, 0, 0 | T2: 1, 0, 0, 0 | T3: 0, 0, 0, 0 | T4: 2, 0, 0, 0 | |
| Gp 25_1 | T1: 1, 0, 0, 0 | T2: 1, 0, 0, 0 | T3: 1, 0, 0, 0 | T4: 2, 0, 1, 0 | b |
| Gp 27_1 | T1: - - - -- | T2: - - - - | T3: - - - - | T4: 2, 0, 0, 0 | |
| Gp 28_1 | T1: 1, 0, 0, 0 | T2: 1, 0, 0, 0 | T3: 1, 0, 0, 0 | T4: 2, 0, 0, 0 | |
| Gp 35_1 | T1: 1, 0, 0, 0 | T2: 1, 0, 0, 0 | T3: 1, 0, 0, 0 | T4: 2, 0, 0, 0 | |
| Gp 38_1 | T1: 0, 1, 0, 0 | T2: 0, 1, 0, 0 | T3: 0, 1, 0, 0 | T4: - - - - | |
| Gp 41_1 | T1: 0, 0, 0, 0 | T2: 0, 0, 0, 0 | T3: 0, 0, 0, 0 | T4: 0, 0, 0, 3 | d |
| Gp 41_2 | T1: 0, 1, 0, 0 | T2: 0, 1, 0, 0 | T3: 0, 0, 0, 0 | T4: 0, 1, 0, 1 | |
| Gp 54_1 | T1: 0, 0, 5, 0 | T2: 1, 0, 3, 0 | T3: 0, 0, 4, 0 | T4: 1, 0, 23, 0 | |
| Gp 54_2 | T1: 0, 0, 0, 0 | T2: 0, 0, 0, 0 | T3: 0, 0, 0, 0 | T4: 3, 0, 15, 3 | b |
| Gp 60_1 | T1: 0, 0, 0, 0 | T2: 0, 0, 0, 0 | T3: 0, 0, 0, 2 | T4: 2 ,0, 0, 4 | |
| Gp 60_2 | T1: 0, 0, 0, 0 | T2: 0, 0, 0, 0 | T3: 0, 0, 0, 2 | T4: 2, 0, 0, 2 | |
| Gp 60_3 | T1: 0, 0, 0, 0 | T2: 0, 0, 0, 0 | T3: - - - - | T4: - - - - | |
| Gp 73_1 | T1: 1, 0, 0, 0 | T2: 0, 0, 0, 2 | T3: 0, 0, 0, 1 | T4: 2, 0, 0, 1 | |
| Gp 73_2 | T1: - - - - | T2: - - - - | T3: - - - - | T4: 20, 0, 0, 11 ensemble | |
| Gp 82_1 | T1: 0, 0, 0, 1 | T2: 1, 0, 0, 0 | T3: 1, 0, 1, 3 | T4: 2, 0, 0 ,14 | |
| Gp 82_2 | T1: 0, 0, 0, 1 | T2: 1, 0, 0, 0 | T3: 1, 0, 0, 4 | T4: 2, 0, 0, 14 | |
| Gp 90_1 | T1: - - - - | T2: - - - - | T3: 1, 0, 0, 0 | T4: 0, 0, 0, 0 | c |
| Gp 91_1 | T1: 1, 0, 0, 0 | T2: - - - - | T3: - - - - | T4: 2, 0, 0, 0 | |

b c d

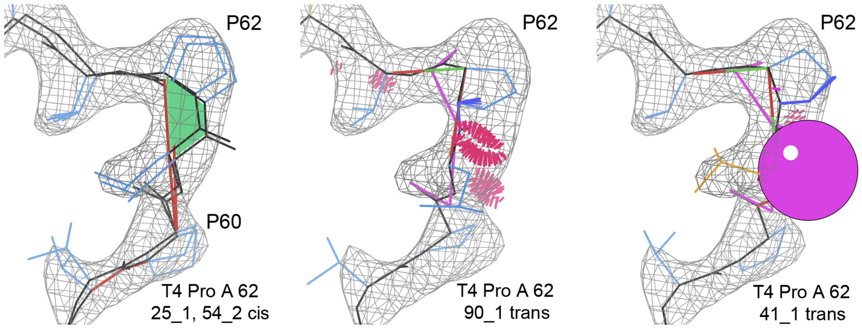

**Supplementary Figure 3. Evaluation of a short sequence misalignment within a helix**
Local Fit-to-Map and Coordinates-only scores are compared for a 3-residue sequence misalignment inside an α-helix in an *ab initio* model submitted to the Challenge (APOF 2.3Å 54_1). (a) Model residues 14-42 vs target map (blue: correctly placed residues, yellow: mis-threaded residues 25-29, black: APOF reference model, 3ajo). (b) Structure-based sequence alignment of the *ab initio* model (top) vs. reference model (bottom). (c) Local Fit-to-Map scores (screenshot from Challenge model evaluation website Fit-to-Map Local Accuracy tool). Curves are shown for Phenix Box_CC (orange), EMDB Atom Inclusion (purple), Q-score (red) EMRinger (green), and SMOC (blue). The score values for model residue Leu 28 are shown in the box at right. (d) Residue scores were calculated using the Molprobity server[2]. The mis-threaded region is boxed in (b-d). Panels (a) and (b) were generated using UCSF Chimera[4].

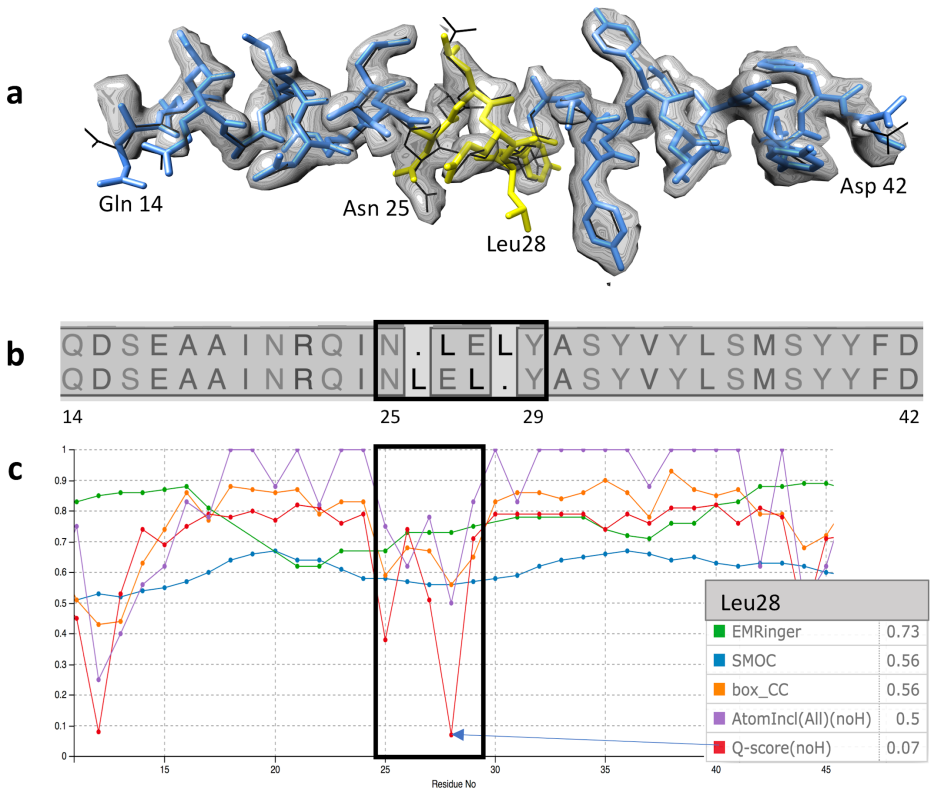


**d**

| Metric | Ile 24 | Asn25 | Leu26 | Glu27 | Leu28 | Tyr29 | Ala 30 |
|---|---|---|---|---|---|---|---|
| Bond Lengths | - | - | Outlier(s) | - | Outlier(s) | Outlier(s) | - |
| Bond Angles | - | Outlier(s) | Outlier(s) | - | Outlier(s) | Outlier(s) | - |
| Clashes | - | - | - | - | Clash | - | - |
| Ramachandran | Favored | Favored | Favored | Allowed | Favored | Favored | Favored |
| CaBLAM | Favored | Disfavored | Disfavored | Favored | Disfavored | Disfavored | Favored |
| Cis Peptides | - | - | - | - | - | CisNonPro | - |

### Supplementary Figure 4. Modeling errors around omitted Zinc ligand in ADH

Target 4 (ADH) density map with examples of modelling errors caused by omission of Zinc ligand. (a) Reference structure with Zinc metal ion (gray ball) coordinated by 4 Cysteine residues (blue sidechains). (b-e) Submitted models missing Zinc (labels indicate the group_model ids). All have geometry and/or conformational violations as flagged by MolProbity CaBLAM (magenta pseudobonds), cis-nonPro (green parallelograms), Ramachandran (green pseudobonds), Cbeta (magenta spheres), and angle (blue and red fans). Model (b) has backbone conformation very close to correct, while (b) and (c) both have flags indicating bad geometry of incorrect disulfide bonds. Models (c) and (d) have backbone distortions, and (e) is mistraced through the Zn density. Molecular graphics were generated using KiNG[3].

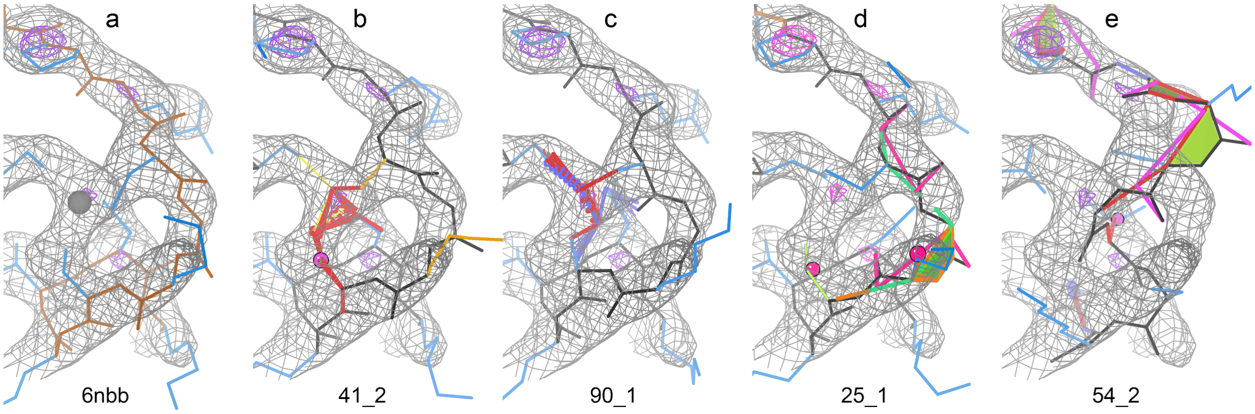


### Supplementary Figure 5. *ab initio* group performance evaluation

Averaged Z-scores are plotted for each modeling group, separated by higher resolution (T1 at 1.8Å, T2 at 2.3 Å) versus lower resolution (T3 at 3.1Å, T4 at 2.9 Å) targets. In each case 6 groups produced very good models (Z ≥ 0.3; green pins), though not the same set of groups (see Group performance section in the main text). Runner-up clusters (-0.3 ≤ Z < 0.3) are shown with gold pins. Individual scores and order shift with alternate choices of evaluation metrics and weights, but the clusters at each resolution level are stable.

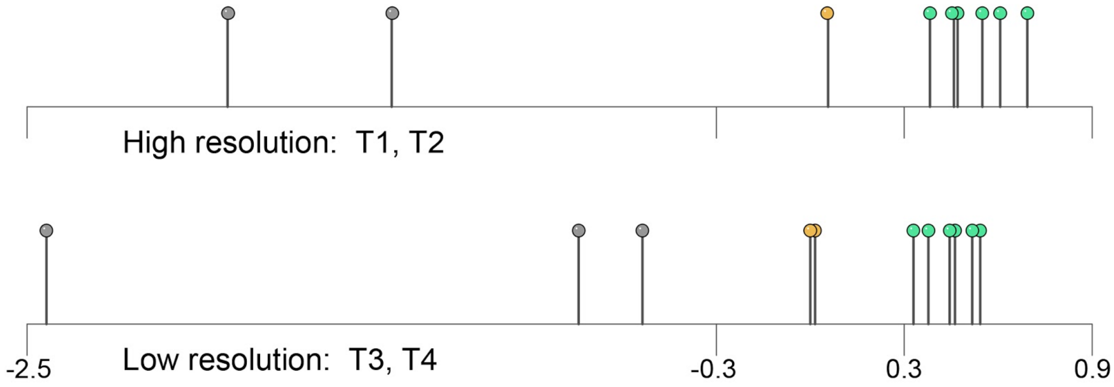
